# Capicua refines mossy fiber–CA3 axon targeting in the late postnatal hippocampus

**DOI:** 10.1101/2025.02.14.638208

**Authors:** Rebekah van Bruggen, Mi Wang, Qiumin Tan

**Author notes:** **Correspondence should be addressed to:** Qiumin Tan, 5-14 Medical Sciences Building, Department of Cell Biology, University of Alberta, Edmonton, Canada T6G 2H7.

## Abstract

Proper brain wiring relies on the precise distribution of axonal projections to specific subcellular domains of their target neurons. These spatially confined connections establish the anatomical foundation for neural circuit assembly. The mossy fiber (MF)–CA3 pathway in the hippocampus is an excellent system to study the mechanisms underlying lamina-specific connectivity. In rodents, MF projections develop postnatally and reach their mature configuration by the end of the second postnatal week. MF axons synapse on the proximal segments of the dendrites but avoid the somas of CA3 pyramidal neurons. As dentate gyrus granule neurons are continuously generated and integrated into the existing hippocampal circuit throughout the postnatal period and adulthood, the mechanisms that guide MF axons to achieve lamina-specific targeting of these later-born granule neurons remain unclear. Here, we show that deletion of the neurodevelopmental disorder-associated protein capicua (CIC) results in abnormal MF targeting in the mouse hippocampus. Notably, this defect emerges after the second postnatal week and persists into adulthood, distinguishing it from classical MF guidance defects, which typically manifest during the first postnatal week. We also demonstrate that this miswiring is due to CIC loss in dentate gyrus granule neurons rather than CA3 pyramidal neurons. Single-nucleus transcriptomics and trajectory analysis reveal a loss of a mature granule neuron subtype and dysregulation of axon guidance genes that are normally downregulated as granule neurons mature. Our findings uncover a previously unrecognized role for CIC in hippocampus development and offer insights into the regulation of lamina-specific MF connectivity in the postnatal brain.

## Introduction

Brain wiring and cognitive functions are defined by synaptic specificity at both cellular and subcellular levels. In many regions of the vertebrate central nervous system, afferent projections are confined to specific laminae enriched for distinct dendritic domains (1–3). The dentate gyrus mossy fiber (MF)–CA3 synapse in the hippocampus is a prime example of a system exhibiting lamina-specific connectivity. MF axons, emanating from dentate gyrus granule neurons, form two distinct bundles: the suprapyramidal and infrapyramidal bundles (**Fig. 1A**). These bundles selectively target the proximal regions of the apical and basal dendrites of CA3 pyramidal neurons, while avoiding the somas located in the stratum pyramidale (4–7). In rodents, MF projections begin forming shortly after birth and achieve their mature configuration by postnatal day 14 (P14) (8). Both repulsive and attractive molecular cues are essential for establishing lamina-specific MF–CA3 connectivity during the first two postnatal weeks (4, 6, 7, 9, 10). However, dentate gyrus granule neurons are continuously generated by progenitor cells in the subgranular zone. These newly born neurons undergo morphological maturation and integrate functionally into the hippocampal network (11, 12). The mechanisms regulating axonal targeting of these later-born granule neurons to maintain MF lamina-specific patterning remain largely unknown. Notably, abnormal MF–CA3 connections have been implicated in learning deficits and epilepsy in animal models (10, 13, 14), underscoring the importance of understanding how lamina-restricted MF targeting is achieved.

**Figure 1.**
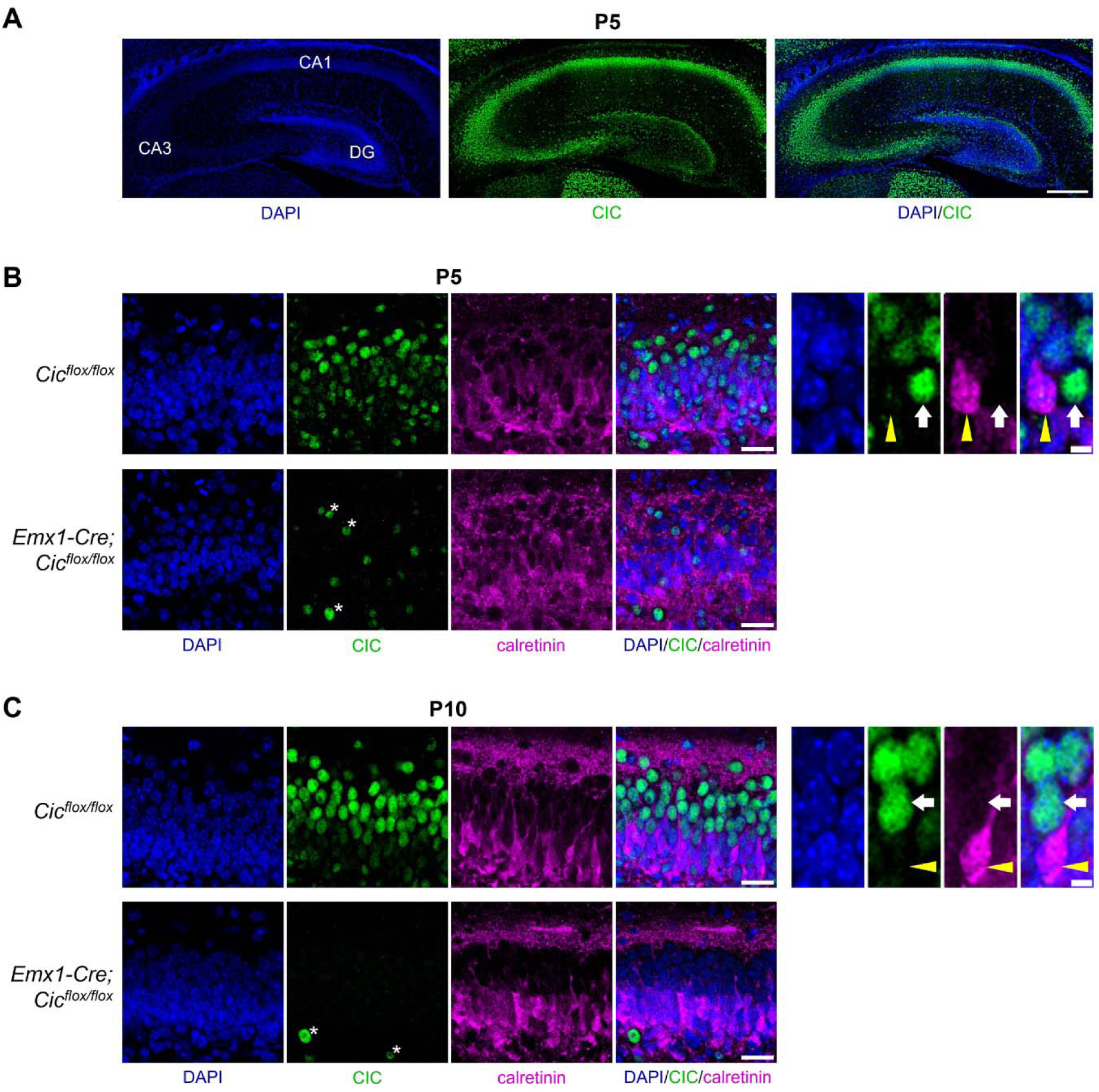
CIC expression in the early postnatal hippocampus. (**A**) Immunofluorescence staining for CIC in the hippocampus at postnatal day (P) 5. CA1, cornu ammonis 1; CA3, cornu ammonis 3; DG, dentate gyrus. Scale bar = 250 µm. (**B**, **C**) Immunofluorescence staining of CIC in the dentate gyrus at P5 (**B**) and P10 (**C**). Calretinin marks immature granule neurons. CIC is strongly expressed in calretinin-negative mature granule neurons (white arrows) and weakly or not expressed in calretinin-positive immature granule neurons (yellow arrowheads). In *Emx1-Cre; Cic^flox/flox^* mice, CIC is efficiently deleted from granule neurons, though CIC immunoreactivity remains detectable (asterisks) in non-*Emx1*-lineage cells, such as inhibitory neurons and microglia. Scale bars = 25 µm for left panels and 5 µm for right panels.

The identification of neurodevelopmental disorder genes often presents unique opportunities to uncover previously unappreciated mechanisms regulating brain development. One such gene is capicua (*CIC*), a transcriptional repressor broadly expressed in the brain (15–17). Heterozygous pathogenic loss-of-function variants in *CIC* result in *CIC*-related neurodevelopmental disorder (MIM #617600), characterized by neurobehavioral phenotypes including learning disabilities, attention-deficit/hyperactivity disorder (ADHD), autism, and epilepsy (15, 18–24). As *CIC*-related neurodevelopmental disorder is rare, mouse models have been developed to study the role of Cic in brain development. While germline homozygous knockout of *Cic* results in perinatal lethality in mice, heterozygous mice show no overt abnormalities (15), suggesting that humans are more sensitive to partial loss of CIC. To better understand the role of CIC in neurodevelopment, conditional knockout mouse models targeting specific brain regions or cell types have been generated (15, 17, 25). For example, deleting *Cic* from forebrain excitatory neurons using the *Emx1-Cre* driver results in ADHD-like behaviors and learning and memory deficits, validating this model as relevant to the human disorder (15). Cellular characterization of these mice has demonstrated the role of CIC in the postnatal maintenance of cortical projection neurons in layers 2–4, with mutant mice exhibiting reduced dendritic complexity and neuron numbers. Moreover, pan-neural *Cic* knockout mice show impaired neuronal differentiation (25), and our recent work has identified a requirement for CIC in dendritic maturation of adult-born hippocampal neurons (26). Collectively, these findings underscore the important role of CIC in neuronal differentiation and dendrite development. However, whether CIC regulates other aspects of neuronal maturation, such as axon guidance and/or targeting, cell adhesion, or synaptogenesis, remains unknown.

In this study, we demonstrate that CIC plays a key role in granule neurons to promote precise MF targeting in the late postnatal mouse hippocampus. Loss of CIC leads to dysregulation of axon guidance genes during granule neuron maturation. Our study provides insights into the role of CIC in hippocampus development and the neurobiology underlying the *CIC*-related neurodevelopmental disorder.

## Results

### Loss of CIC leads to abnormal mossy fiber spreading to the CA3 stratum pyramidale region

To assess the role of CIC in the developing hippocampus, we first examined its expression pattern and found that, at P5, CIC was expressed in CA1, CA3, and dentate gyrus (**Fig. 1A**). CIC expression appeared to be ubiquitous throughout the CA1 and CA3 but was most concentrated in cells at the outer edge of the granular layer of the dentate gyrus, where mature granule neurons reside. To determine whether CIC expression correlates with neuronal maturation, we used calretinin as a marker for immature neurons (27). At P5, CIC was highly expressed in calretinin-negative (mature) granule neurons, while its expression was low to undetectable in calretinin-positive (immature) neurons (**Fig. 1B**). This pattern became even more pronounced at P10, as the spatial segregation of immature and mature granule neurons within the granular layer became more distinct (**Fig. 1C**). The upregulation of CIC from immature to mature neurons indicates a role for CIC in neuronal maturation in the developing dentate gyrus.

To further explore the role of CIC in the early postnatal hippocampus, we utilized the *Emx1-Cre; Cic^flox/flox^* mice, in which Cre-mediated recombination begins in neural progenitors at embryonic day 9.5 (15, 26, 28). Consequently, CIC is deleted from dentate gyrus granule neurons and CA neurons in the hippocampus (**Figs. 1B, C**, and **S1**). In rodents, MF axons develop from P0 to P14 (8) and bifurcate into the suprapyramidal and infrapyramidal bundles that synapse on either side of the CA3 stratum pyramidale (SP), where CA3 neuron cell bodies reside (**Fig. 2A**). To investigate the impact of CIC deletion on hippocampal development, we compared the MF pathway in control (*Cic^flox/flox^*) and *Emx1-Cre; Cic^flox/flox^* knockout mice using calbindin (CALB1) immunostaining, a common marker for mature granule neurons and their MF projections (4, 6, 9, 29). At P20, when MF pathfinding is complete, control mice exhibited a lamina-specific projection of the MF suprapyramidal bundle, which primarily innervated the stratum lucidum (SL), where the apical dendrites of CA3 pyramidal neurons are located, with few CALB1^+^ MF terminals found in the SP (**Fig. 2B**–**D**). In knockout mice, this pattern was altered. Although the suprapyramidal bundle still projected to the SL, numerous CALB1^+^ MF terminals abnormally spread into the SP (**Fig. 2B–D**). These aberrant MF terminals co-expressed the zinc transporter ZnT3 (**Fig. 2E**), typically localized to MF axon terminals (29–31), as well as the pre-synaptic marker synaptophysin (**Fig. 2F**). Our quantifications further revealed a significant increase in both the number (**Fig. 2G**) and size (**Fig. 2H**) of the CALB1^+^ ZnT3^+^ MF terminals in the knockout SP compared to controls. Additionally, immunostaining and immunoblotting for synapsin III (SYN3), a protein highly enriched in MF synapses (32, 33), demonstrated increased SYN3 levels in the SP of knockout mice (**Fig. 2I, J**).

**Figure 2.**
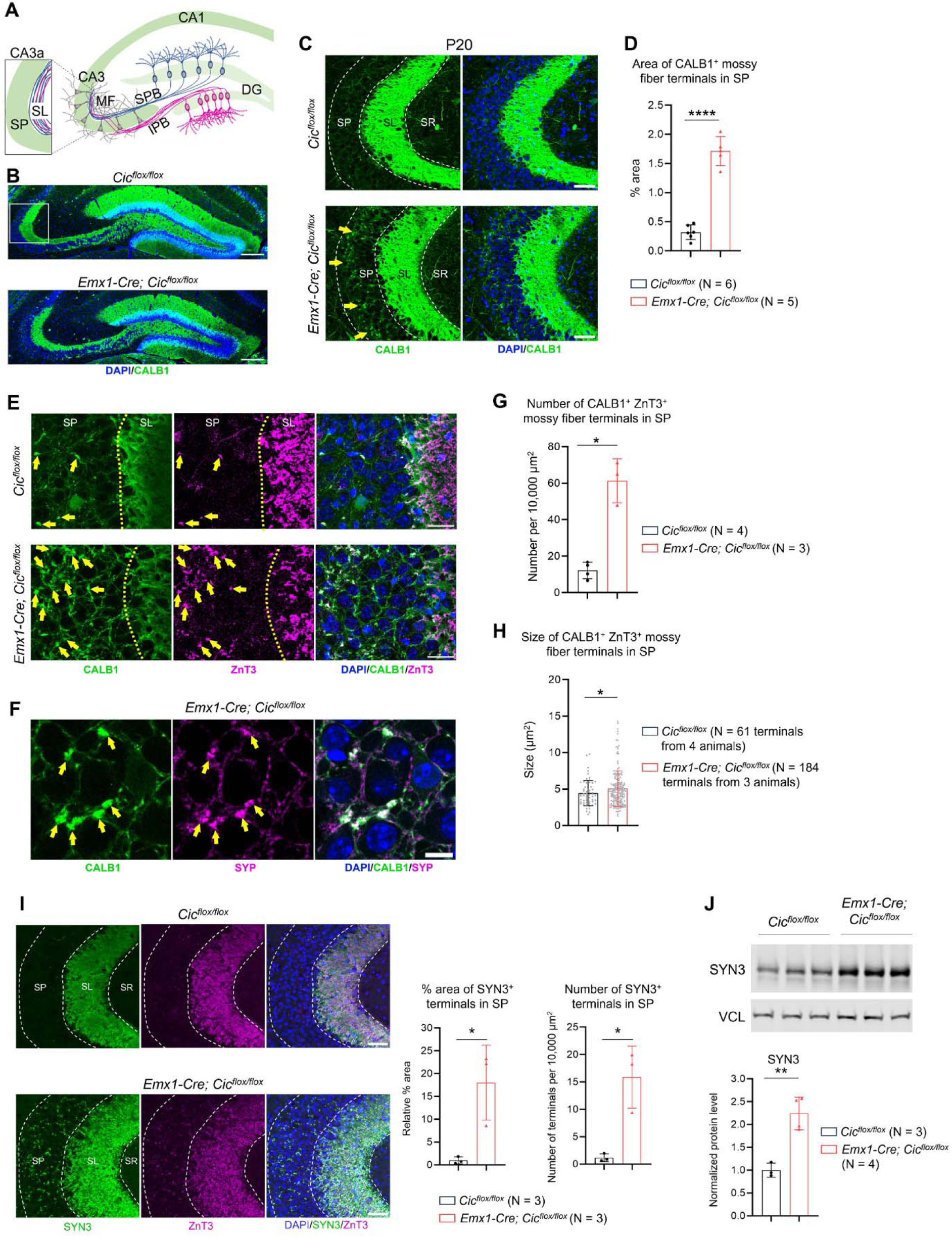
Abnormal mossy fiber targeting in *Emx1-Cre; Cic^flox/flox^* knockout mice. (**A**) Schematic of the mouse hippocampus illustrating the mossy fiber (MF) pathway. DG, dentate gyrus; SPB, suprapyramidal bundle; IPB, infrapyramidal bundle; SL, stratum lucidum; SP, stratum pyramidale. (**B**) Representative confocal images of calbindin (CALB1) immunostaining in coronal sections of the hippocampus from postnatal day (P) 20 mice. Scale bars = 200 µm. (**C**) The CA3a area (white box in **B**) is shown at a higher magnification. Dotted lines demarcate the SP and SL of CA3a. Arrows indicate aberrant MF invasion into the SP in knockout mice. SR, stratum radiatum. Scale bars = 50 µm. (**D**) Quantification of CALB1^+^ immunoreactive areas in the SP. (**E**) Co-localization of CALB1 and zinc transporter 3 (ZnT3) (arrows) confirms the presence of abnormal MF axon terminals in the SP in knockout mice. Scale bars = 25 µm. (**F**) Co-localization of CALB1 and synaptophysin (SYP) (arrows) identifies presynaptic terminals in the SP. Scale bar = 10 µm. (**G** and **H**) Quantification of the number (**G**) and size (**H**) of CALB1^+^ ZnT3^+^ MF terminals in the SP. (**I**) Abnormal MF terminals in knockout mice express synapsin III (SYN3), a marker of MF synapses. Quantifications of the area and number of SYN3^+^ MF terminals are shown on the right. Scale bars = 50 µm. (**J**) Immunoblot analysis and quantification of SYN3 levels in the knockout DG at P19 confirming SYN3 upregulation. Data are presented in scatter plots with error bars representing ± SD. Statistical analysis was performed using Welch’s t test. *, *P* < 0.05; **, *P* < 0.01; ****, *P* < 0.0001.

Next, we investigated when the abnormal MF innervation becomes evident in the *Emx1-Cre; Cic^flox/flox^* knockout mice. At P11, CALB1 immunostaining revealed similar MF projections between control and knockout mice (**Fig. 3A**). By P14, however, a mild but significant increase in MF terminals was observed in the SP of knockout mice, although the difference was not as pronounced as at P20, indicating that abnormal MF targeting becomes more prominent during the third postnatal week (**Fig. 3B**). This abnormal MF spreading peaks around 11 weeks of age and appears to stabilize thereafter till at least one year of age. Altogether, our data demonstrate that ectopic MF terminals begin forming in the CA3 SP of *Emx1-Cre; Cic^flox/flox^* knockout mice around the second postnatal week and persist throughout adulthood.

**Figure 3.**
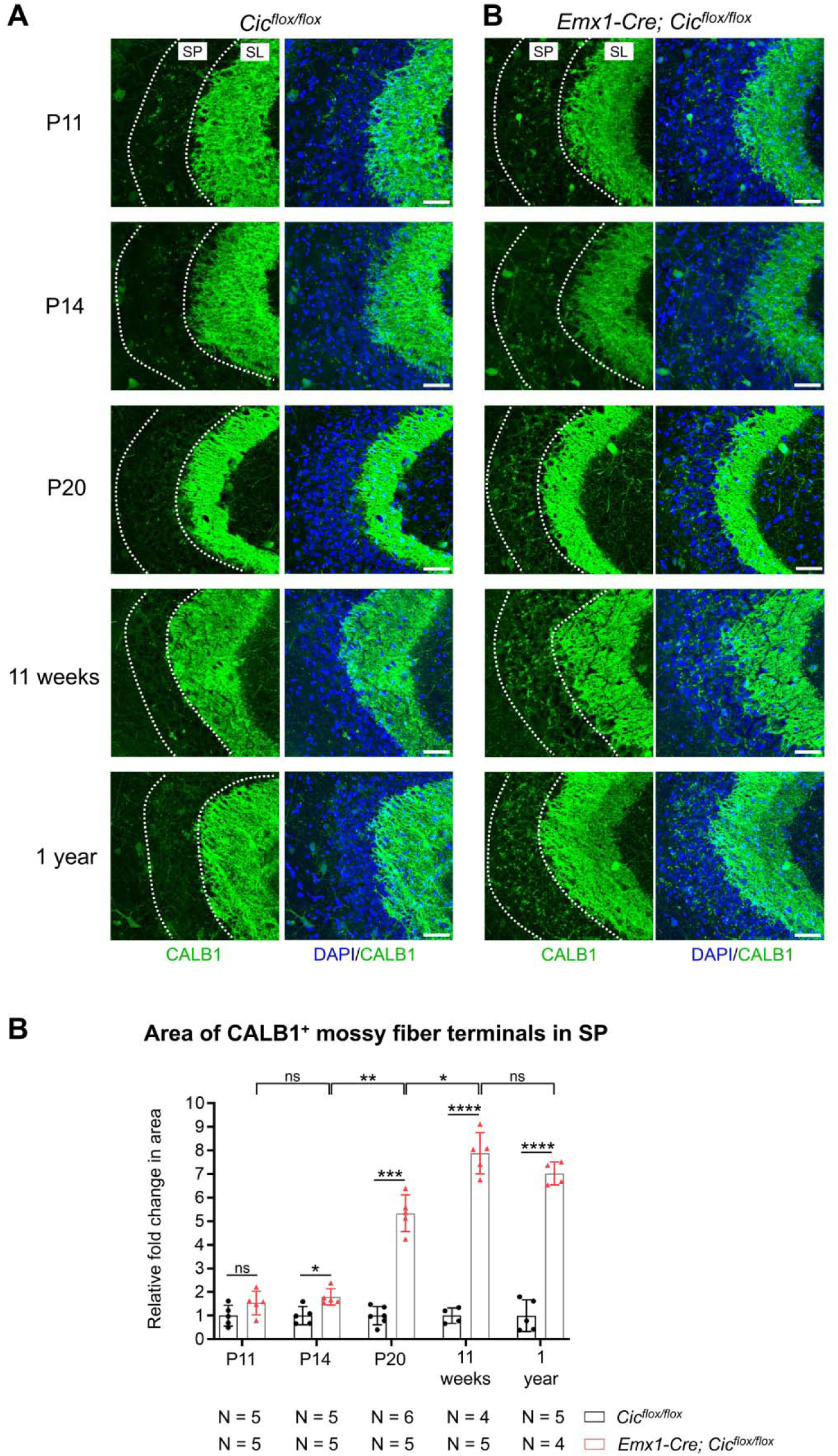
Persistent aberrant mossy fiber–CA3 innervation in *Emx1-Cre; Cic^flox/flox^* knockout mice. (**A**) Representative confocal images from postnatal day (P) 11, P14, P20, 11 weeks, and 1-year-old mice, showing abnormal calbindin (CALB1)-positive mossy fiber terminals in the stratum pyramidale (SP) of the CA3a region in *Emx1-Cre; Cic^flox/flox^* knockout mice. SL, stratum lucidum. Scale bars = 50 µm. (**B**) Quantification of CALB1^+^ immunoreactive area within the SP across different ages. Data are presented as a scatter plot with error bars indicating mean ± SD. Statistical analysis was performed using mixed-model two-way ANOVA, followed by Holm-Sidak *post hoc* test for multiple comparisons. ns, not significant; *, *P* < 0.05; **, *P* < 0.01; ***, *P* < 0.001; ****, *P* < 0.0001.

### CIC is required in dentate gyrus granule neurons to promote lamina-specific mossy fiber innervation

Since CIC is deleted from both dentate gyrus granule neurons and CA3 pyramidal neurons in the *Emx1-Cre; Cic^flox/flox^* mice (**Fig. S1**), the abnormal MF terminals observed in the CA3 SP of these mice could be due to CIC loss in either neuron type. To distinguish between these possibilities, we utilized the *Rbp4-Cre* driver line to selectively delete CIC from dentate gyrus granule neurons (4, 34). Using a Cre-dependent tdTomato (tdT) reporter line, we validated that Cre-mediated recombination was restricted to granule neurons at P20, with no tdT-expressing CA3 neurons detected (**Fig. 4A**). Next, we generated *Rbp4-Cre; Cic^flox/flox^; tdT* knockout mice and confirmed efficient CIC deletion in 92.1±7.2% (N = 3 animals) of tdT-expressing granule neurons at P20 (**Fig. 4B**). The expression of tdT in granule neurons enabled us to compare the MF pathway in control versus knockout mice. At P20, tdT^+^ MF projections in control mice were predominantly confined to the SL of the CA3, with minimal terminals in the SP, resembling the pattern of CALB1 expression (**Fig. 4C**). In contrast, tdT^+^ MF projections extended into the SP in the *Rbp4-Cre; Cic^flox/flox^* knockout mice, and these abnormally projecting MF terminals co-expressed CALB1 and synaptophysin (**Fig. 4C-E**). Ectopic MF projections persisted in adult knockout mice at 8 weeks of age (**Fig. S2**). Interestingly, the relative area of SP occupied by aberrant MF terminals was smaller in the *Rbp4-Cre; Cic^flox/flox^* mice compared to the *Emx1-Cre; Cic^flox/flox^* mice (comparing **Figs. 3B**, **4D**, and **S2B**). This may be because CIC was deleted from nearly all granule neurons in the *Emx1-Cre; Cic^flox/flox^* mice (**Fig. 1**), whereas only a subset of granule neurons was affected in the *Rbp4-Cre; Cic^flox/flox^* mice (**Fig. 4B**). Overall, our data demonstrate that CIC deletion in dentate gyrus granule neurons alone is sufficient to induce persistent ectopic MF innervation to the SP.

**Figure 4.**
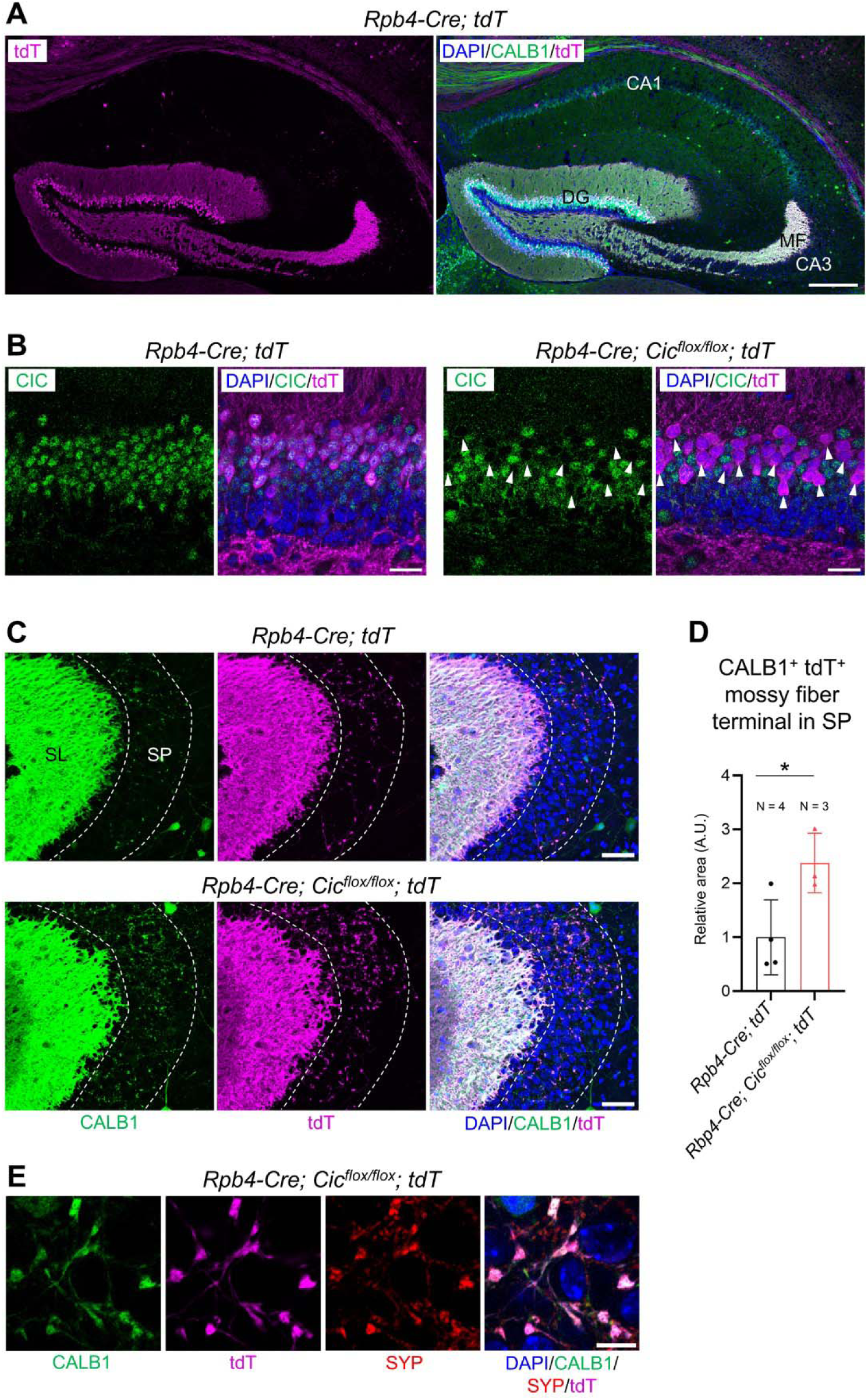
CIC is required in dentate gyrus granule neurons to promote lamina-specific mossy fiber innervation. (**A**) Confocal images of the hippocampus in *Rbp4-Cre; tdT* mice at postnatal day (P) 20. Scale bar = 250 µm. CA1, cornu ammonis 1; CA3, cornu ammonis 3; MF, mossy fiber. (**B**) Efficient CIC deletion in tdT^+^ granule neurons (arrowheads) is observed in *Rbp4-Cre; Cic^flox/flox^; tdT* mice. Scale bars = 25 µm. (**C**) In *Rbp4-Cre; Cic^flox/flox^; tdT* mice, mossy fiber (MF) axon terminals, marked by CALB1 and tdT, abnormally innervate the stratum pyramidale (SP) of the CA3a region. SL, stratum lucidum. Scale bars = 50 µm. (**D**) Quantification of CALB^+^ TdT^+^ MF terminal area in the SP. Data are presented as a scatter plot with error bars indicating mean ± SD. Statistical analysis was performed using Welch’s t test. *, *P* < 0.05. (**E**) Co-localization of CALB1, tdT, and synaptophysin (SYP) identifies presynaptic MF terminals in the SP. Scale bar = 10 µm.

### Loss of CIC in CA3 neurons does not affect mossy fiber projections

To investigate whether CIC is also required in CA3 neurons for precise MF targeting, we initially attempted to delete CIC from CA3 neurons using the *Grik4-Cre* line (35). Incorporating the Cre-dependent *tdT* allele, we observed extensive Cre-mediated reporter recombination not only in CA3 neurons but also in dentate gyrus granule neurons by P20 (**Fig. S3A**), consistent with previous reports (35, 36). However, Cre activity was insufficient to mediate recombination of the *Cic^flox/flox^* alleles, as no CIC deletion was detected in tdT^+^ granule neurons in *Grik4-Cre; Cic^flox/flox^* mice (**Fig. S3B**). On the other hand, we observed CIC loss in a modest (∼25%) number of CA3 neurons (**Fig. S3C**). Therefore, CIC deletion in the *Grik4-Cre; Cic^flox/flox^* mice was restricted to CA3 neurons. Analysis of the MF pathway using CALB1 immunostaining revealed no differences between control and knockout mice at P20 (**Fig. S3D**), suggesting that partial loss of CIC in CA3 neurons does not affect lamina-restricted MF targeting.

To further assess the role of CIC in CA3 neurons, we employed a viral approach to delete *Cic* in these cells. We performed intracerebroventricular injections of adeno-associated virus (AAV) carrying a Cre recombinase and tdT reporter under a neuron-specific promoter (AAV8/*hSyn-Cre-P2A-tdTomato* or AAV8/*Cre* for short) into *Cic^flox/flox^* pups at P0 (**Fig. S4A**). Due to the high proliferative activity of granule neuron precursors at P0, this approach more efficiently targets post-mitotic CA3 neurons (37, 38). By adjusting the viral titer, we generated a mosaic system that allows for the transduction of only a small subset of granule neurons and some, but not all, CA3 neurons. In control animals (*Cic^flox/+^* pups injected with AAV8/*Cre*), CIC expression remained detectable in AAV-transduced CA3 neurons, marked by tdT expression (**Fig. S4B**, yellow arrowheads). In contrast, CIC immunoreactivity was absent in tdT^+^ CA3 neurons of *Cic^flox/flox^* pups injected with AAV8/*Cre* (**Fig. S4B**, white arrowheads), confirming efficient gene deletion. This mosaic system enabled us to determine whether the soma of a single transduced CA3 neuron was aberrantly surrounded by MF terminals. If CIC is required in CA3 neurons for proper MF projections, we would expect wildtype MF axons to innervate CIC-deleted, tdT-expressing CA3 neuron somas. However, this was not observed in any of the AAV-transduced, CIC-deleted CA3 neurons (**Fig. S4C**). Collectively, these results indicate that CIC is not required in CA3 neurons to maintain lamina-specific MF targeting.

### Dysregulation of axon guidance genes during granule neuron maturation in the *Emx1-Cre; Cic^flox/flox^* knockout mice

To gain insights into the molecular mechanism by which CIC sculpts the MF pathway, we performed single-nucleus RNA sequencing (snRNA-seq) on hippocampal tissue from P19 control (*Cic^flox/flox^*) and *Emx1-Cre; Cic^flox/flox^* knockout mice. After quality control, we retained 17,749 nuclei from the control dataset, including 5,871 granule neuron lineage cells (identified by *Prox1* expression), and 16,871 nuclei from the knockout dataset, consisting of 5,627 granule neuron lineage cells. We identified seven granule neuron clusters (**Fig. 5A, Fig. S5A–C, Table S1**), categorized into three immature (imGN1–3) and four mature (mGN1–4) subsets based on their expression of the immature neuron marker *Dcx* and the mature neuron marker *Calb1* (**Fig. 5B**). Gene ontology (GO) analysis of cluster markers revealed that imGN1 was enriched for terms related to transcription, imGN2 and imGN3 for axon development, and mGN1–4 for synaptic transmission, supporting their maturation stages (**Table S2**). Strikingly, we found a dramatic reduction in the mGN4 cluster in knockout mice, while the cell numbers in other clusters were comparable (**Fig. 5C, D**). The mGN4 cluster uniquely expressed *Ntng1* (nectrin G1) and *Itgav* (integrin alpha-V) (**Fig. 5E**), both with spatial and temporal expression patterns consistent with a mature granule neuron subtype (39) (**Fig. S5D**).

**Figure 5.**
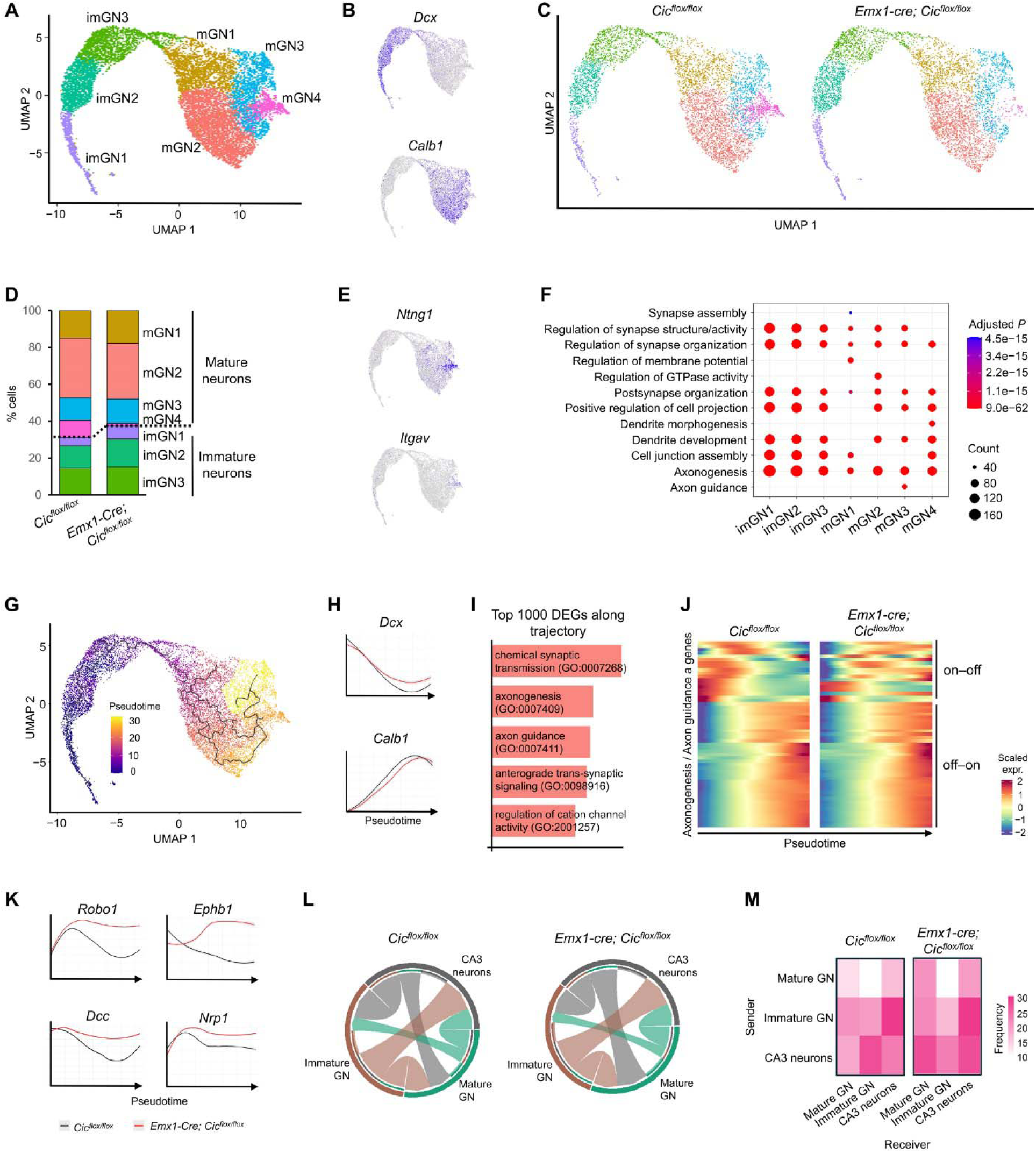
Dysregulation of axonogenesis genes during granule neuron maturation in *Emx1-Cre; Cic^flox/flox^* knockout mice. (**A**) 2D uniform manifold approximation and projection (UMAP) plot showing seven clusters of granule neurons, including immature (imGN) and mature (mGN) granule neurons. (**B**) UMAP feature plots illustrating the expression of *Dcx* and *Calb1*, markers for immature and mature granule neurons, respectively. (**C**) UMAP plots comparing the distribution of granule neuron clusters in control (*Cic^flox/flox^*) versus knockout (*Emx1-Cre; Cic^flox/flox^*) mice. (**D**) A bar plot showing the proportion of cells within each cluster across the two datasets. (**E**) UMAP feature plots revealing the expression of *Ntng1* and *Itgav* within the mature neuron cluster mGN4. (**F**) A dot plot highlighting the top seven gene ontology (GO) terms for differentially expressed genes (DEGs) across all clusters. (**G**) UMAP plot depicting the maturation trajectory of granule neurons, with cells colored according to pseudotime values. (**H**) Pseudotime expression plots showing the gene expression profiles of *Dcx* and *Calb1* across the granule neuron maturation trajectory. (**I**) The top five GO terms associated with the top 1,000 DEGs along the granule neuron trajectory in control mice. (**J**) Heatmap analysis of axonogenesis and axon guidance genes across pseudotime. “On–off” genes refer to those with high expression in immature granule neurons and low expression in mature neurons, while “off–on” genes are highly expressed only in mature granule neurons. (**K**) Pseudotime expression plots for *Robo1*, *Ephb1*, *Dcc*, and *Nrp1*, revealing their expression dynamics throughout granule neuron maturation. (**L**) Frequency chord diagrams illustrating overall cell–cell communications among different cell populations. Each cell population is depicted as a segment along the outer ring of the circular layout, with the size of the arcs between segments proportional to the frequency of interactions between potentially communicating cell types. (**M**) Heatmaps showing the frequency of cell–cell communication among various cell populations.

We then compared gene expression between the control and knockout datasets and examined pathways associated with differentially expressed genes (DEGs). In *Emx1-Cre; Cic^flox/flox^* knockout mice, we observed upregulation of CIC target genes *Etv1*, *Etv4*, and *Etv5* (40) across all seven clusters, validating the knockout model and sequencing approach (**Fig. S6**). Gene ontology (GO) analysis revealed that “axonogenesis” was the most enriched term among DEGs across all clusters (**Fig. 5F**), despite axon-related terms not being enriched in mature granule neuron subsets (**Table S2**). This led us to hypothesize that axonogenesis genes may be upregulated in mature granule neurons in the knockout mice. To test this, we performed pseudotime analysis of gene expression dynamics along the granule neuron maturation trajectory (**Fig. 5G**). This analysis confirmed *Dcx* expression decreased while *Calb1* expression increased along pseudotime in both control and knockout mice, indicating a similar overall maturation trajectory (**Fig. 5H**). Go analysis of the top 1000 DEGs along the granule neuron trajectory in control mice showed enrichment for axon- and synapse-related terms (**Fig. 5I**). Heatmap analysis of axonogenesis-related genes in control mice revealed two patterns: genes highly expressed in immature neurons but downregulated in mature neurons (“on–off” genes), and genes upregulated during granule neuron maturation (“off–on” genes) (**Fig. 5J**). In knockout mice, “off–on” gene expression was preserved, but the dynamics of “on–off” genes were specifically disrupted, with many being expressed later in pseudotime and remaining elevated throughout maturation. These genes included key axon guidance receptors such as *Robo1*, *Ephb1*, *Dcc*, and *Nrp1* (**Fig. 5K**). We next asked whether the upregulation of axon-related genes affected interactions between granule neurons and their downstream target CA3 neurons. To this end, we re-clustered the hippocampus snRNA-seq data to include CA3 pyramidal neurons alongside immature (imGN1–3) and mature (mGN1–4) granule neurons. Using a cell–cell communication analysis framework (41), we found that CA3 neurons exhibited increased interaction with mature granule neurons and decreased interaction with immature granule neurons in the knockout mice (**Fig. 5L, M**). Overall, our findings suggest that CIC loss impedes the development of a specific mature granule neuron cluster and disrupts the expression dynamics of axon-related genes, particularly those with highest expression in immature granule neurons.

## Discussion

Significant headway has been made in defining the mechanisms that initially establish the lamina-specific MF pathway in the early postnatal hippocampus. But what regulates MF pathfinding in later-born granule neurons remains elusive. Here, we demonstrate that loss of CIC in dentate gyrus granule neurons results in aberrant MF terminals in the CA3. This defect is associated with dysregulation of a distinct set of axonogenesis genes during granule neuron maturation. Our findings not only provide insights into the poorly understood process of lamina-specific MF–CA3 circuit assembly in the juvenile and adult hippocampus but also shed light on the neurobiology of *CIC*-related neurodevelopmental disorder.

In the developing hippocampus, MF axons first innervate the CA3 around P0, with their lamina-restricted distribution established by the end of the second postnatal week (8). This early developmental pathfinding is primarily guided by repulsive interactions between class A plexins and their semaphorin ligands (6, 9, 10). Complete loss of semaphorin or plexin function results in global disruption of MF targeting, typically evident by P5–P10 (6, 9). In contrast, the MF mis-targeting phenotype in *Cic* knockout mice emerges after P14 without affecting the overall MF pathway, suggesting a distinct mechanism beyond semaphorin/plexin-mediated repulsion. Indeed, aside from *Nrp1*, an essential co-receptor for class 3 semaphorin signaling (42–44), we did not observe prominent dysregulation of other semaphorin or plexin family members. This suggests that semaphorin–plexin signaling establishes the initial “blueprint” for MF pathfinding during the first two postnatal weeks, while a CIC-dependent mechanism refines MF targeting at later stages. This notion aligns with the ongoing neurogenesis of granule neurons, where new neurons born from progenitors integrate into existing hippocampal circuits, with MF targeting potentially relying on different cues in juvenile and adult-born neurons compared to perinatally generated ones (11, 12). Given that *CIC* is more highly expressed in mature granule neurons, it may play a role in refining MF terminals as neurons mature. Notably, MF axon terminals continuously remodel in response to experience and aging (45), raising the possibility that *CIC* may be involved in maintaining proper MF–CA3 connectivity throughout life.

We observe that CIC level is low in immature granule neurons but upregulated in mature ones, a pattern that persists from early postnatal development into adulthood (26). The increasing prevalence of MF mis-targeting in *Cic* knockout mice over time suggests a continuous role for CIC in maintaining precise MF targeting. GO analysis of cluster markers indicates a functional shift from an axonogenesis program in immature granule neurons to neurotransmission in mature neurons. Upregulation of CIC in mature granule neurons may be necessary for this transition. Pseudotime analysis supports this idea, demonstrating that in the absence of CIC, this switch is impaired, leading to the persistent expression of axonogenesis genes. Strikingly, only axonogenesis genes with an “on–off” expression pattern are disrupted, while those with an “off– on” pattern are spared, suggesting CIC plays a role in repressing specific genes as granule neurons mature. The precise contribution of these dysregulated genes to the MF mis-targeting phenotype remains unclear. Some, like *Robo1*, *Ephb1*, *Dcc*, and *Nrp1*, are well-known for their roles in axon guidance (3, 46–51). Of particular interest is *Ephb1*, which mediates reverse signaling in CA3 neurons to control MF pruning and targeting (29). How upregulation of *Ephb1* in mature granule neurons in the *Cic* knockout mice affects these processes requires further investigation. Future studies should aim to delineate the exact roles of these receptors, with a focus on their localization in granule neuron lineage cells and the signaling pathways involved.

We also uncovered a drastic reduction in a subset of mature granule neurons expressing *Ntng1* in the *Cic* knockout mice at P19, the time of our snRNA-seq analysis. Previous research indicates that *Ntng1*-positive cells are sparse before P16 but increase significantly by P24 and become widely distributed throughout the granular layer by P37 (39). The *Ntng1*-positive cell cluster we identified likely represents an emerging subset of mature granule neurons that later become more prevalent in the dentate gyrus. The function of these *Ntng1*-expressing mature granule neurons remains to be determined. Interestingly, NTNG1 and its ligand NGL-1 have been shown to regulate lamina-specific subdendritic segments in the entorhinal cortex to dentate gyrus pathway, whereby entorhinal cortex axons express NTNG1 while granule neuron dendrites express NGL-1 (52). Our data, together with data from others (39), raise the possibility that *Ntng1*-expressing granule neurons may regulate lamina-specific MF targeting in a similar fashion. Nonetheless, the reduction in *Ntng1*-expressing granule neurons in *Cic* knockout mice suggests a block or delay in neuronal maturation. Whether this defect resolves later in adulthood and how it contributes to the MF mis-targeting phenotype remains to be explored.

Individuals with the *CIC*-related NDD exhibit learning disabilities, and ∼30–40% of them also have epilepsy (15, 22, 53, 54). Given the essential role of the hippocampus in learning, memory, and seizure regulation, disruption of MF–CA3 connections and the loss of *Ntng1*-expressing granule neurons may impair hippocampal function, contributing to the cognitive and epileptic phenotypes seen in these individuals. Our findings therefore not only lend new insight into hippocampus development and brain wiring, but also identify key molecular and cellular pathways that may contribute to the pathophysiology of CIC-related NDD.

## Materials and Methods

### Sex as a biological variable

Our study examined male and female animals, and similar findings are reported for both sexes.

#### Mice

*Emx1-Cre* mice [B6.129S2-*Emx1^tm1(cre)Krj^*/J, #005628], *Grik4-Cre* mice [C57BL/6-Tg(Grik4-cre)G32-4Stl/J, #006474], and Ai9 mice [B6.Cg-*Gt(ROSA)26Sor^tm9(CAG-tdTomato)Hze^*/J, #007909] were obtained from The Jackson Laboratory. Generation of the *Cic^flox^* mice was previously described (15); the *Cic^flox^* mice were also available from The Jackson Laboratory (#030555). Wildtype mice were originally obtained from The Jackson Laboratory (#000664) and subsequently bred in-house. *Rbp4-Cre* mice [STOCK Tg(Rbp4-cre)KL100Gsat/Mmucd, # 031125-UCD] were obtained from The Mutant Mouse Resource and Research Center. Mice were group-housed in a 12-hour light – 12-hour dark cycle, with all experiments performed during the light period. Ages are indicated where applicable. The experimenters were blinded to the genotypes of the mice during the experiment and/or when assessing the outcome. All procedures in mice were approved by the Animal Care and Use Committee of the University of Alberta (AUP 2665 and AUP 3930). All methods were performed in accordance with the relevant guidelines and regulations.

#### Tissue preparations

Mice were deeply anesthetized using i.p. injection of sodium pentobarbital (Euthanyl, 240 mg/mL, Bimeda-MTC, Cambridge, ON), then transcardially perfused with phosphate buffered saline (PBS, Fisher Bioreagents, Cat# BP399-20) followed by 4% paraformaldehyde (Electron Microscope Sciences, Cat# 19202) in PBS. The brains were dissected and further immersed in 4% paraformaldehyde overnight at 4°C, followed by sequential submersion in 15% and then 30% sucrose in PBS, each for 24 hours at 4°C. Brain tissues were cut using a coronal brain matrix, embedded into Optimal Cutting Temperature (OCT) compound (Fisher HealthCare, Cat# 4585) and frozen at −80°C. Coronal brain sections (40 µm-thick) were cut using a cryostat (Leica Microsystems Inc., Cat# CM1520) and kept at 4°C in PBS with 0.02% sodium azide (BICCA, Cat# 7144.8-16). When desired, coronal sections were transferred onto Superfrost Plus Microscope slides (Fisher Scientific, Cat# 12-550-5) and air-dried overnight. Once the slides were dry, they were used immediately for immunofluorescence staining or stored at −80°C.

#### Immunohistochemistry

Slides were post-fixed in 10% phosphate buffered formalin (Fisher Chemicals, Cat# SF100-4) for 10 min at room temperature and then washed in PBS. Antigen retrieval was performed with citric-acid based antigen unmasking solution (VectorLabs, Cat# H-3300) for 30 minutes at 95°C in a water bath. Once the slides were cooled to room temperature, they were washed twice with PBS, permeabilized with PBST (PBS + 0.3% Triton X-100, Fisher Bioreagents, Cat# BP151-500) for 20 min at room temperature and then blocked with 5% normal donkey serum (Sigma, Cat# D9663-10ML) diluted in PBST (blocking solution) for 20 min at room temperature. Primary antibodies were diluted in blocking solution, added onto the slides, and incubated overnight at 4°C in a humid chamber. Sections were washed three times with PBST prior to incubating for 2 hours at room temperature in secondary antibody diluted in blocking buffer. Afterwards, the slides were washed in PBST then PBS and autofluorescence was quenched using Vector TrueView autofluorescence quenching kit, prepared as per manufacturer’s instructions (VectorLabs, Cat# SP8400). The slides were washed, counterstained with DAPI (5 µg/mL, Invitrogen, Cat# D3571) for a 10-min incubation at room temperature, then washed with PBS. The slides were then mounted using VectaShield Vibrance antifade mounting media (VectorLabs, Cat# H170010) and covered with a cover slip. The slides were left to dry overnight, sealed with transparent nail polish, then further dried prior to being imaged with a confocal microscope. The following primary antibodies were used: goat anti-tdTomato (SICGEN Cat# AB8181-200, RRID:AB_2722750, 1:500), rabbit anti-CALB1 (Swant Cat# CB38, RRID:AB_10000340, 1:500), mouse anti-ZnT3 (Synaptic Systems Cat# 197011, RRID:AB_2189665, 1:500); rabbit anti-CIC ((15); RRID:AB_2721281; 1:500); guinea pig anti-SYP (Synaptic Systems Cat# 101308, RRID:AB_2924959; 1:500); rabbit anti-SYN3 (Synaptic Systems Cat#106303, RRID:AB_2619775, 1:500). The secondary antibodies used were donkey anti-rabbit AlexaFluor488 (Invitrogen Cat# A21206, RRID:AB_2535792, 1:1000), donkey anti-goat AlexaFluor555 (Invitrogen Cat# A21432, RRID:AB_2535853, 1:1000), donkey anti-mouse AlexaFluor647 (Invitrogen Cat# A31571, RRID:AB_162542, 1:1000), donkey anti-guinea pig AlexaFluor647 (Jackson ImmunoResearch Labs Cat# 706-605-148, RRID:AB_2340476, 1:1000).

#### Adeno-associated viruses and neonatal intracerebroventricular injections

*pAAV-hSyn-Cre-P2A-dTomato* was a gift from Rylan Larsen (Addgene viral prep # 107738-AAV8; http://n2t.net/addgene:107738; RRID:Addgene_107738). Viruses were aliquoted and stored at −80°C until use. Thawed viruses were kept at 4°C and used within a week. All viruses were diluted to 1.0 X 10^12^ genome copy (GC)/mL using PBS before injection into mice. Neonatal intracerebroventricular injections were performed as previously described (55). Briefly, newborn pups (within 6 hours of birth) and dam were transported to the surgery suite in their home cage. Half of the litter was removed from the cage and transferred to a biological safety cabinet. Each pup was anesthetized using hypothermia and the cranial surface was disinfected with a 70% ethanol wipe (Becton Dickinson Canada Inc, Cat# 326910). Using a gas-tight syringe (Hamilton, Cat# 361025642) with a 32G 1.25-cm needle (Hamilton, Cat# 7762-03), 2 µL of AAV was injected into the lateral ventricles of each hemisphere by hand. The pup was then placed on a 37°C heating pad until recovered and the remaining half of the litter was completed. The injected pups were then returned to the home cage, followed by removing the second half to the litter for AAV injection.

#### Protein extraction and immunoblotting

Tissues (one dentate gyrus per extraction) were homogenized using Dounce Tissue Grinders (DWK Life Sciences, Cat# 8853000002) and 300 µL of cold T-PER Tissue Protein Extraction Reagent (Thermo Fisher Scientific, Cat# 78510) supplemented with fresh protease and phosphatase inhibitors (Thermo Fisher Scientific, Cat# A32953 and Cat# A32957, respectively). Lysates were incubated on ice for 10 min and then cleared by centrifugation at 16,000 xg for 10 min at 4°C. Protein concentrations were quantified using the BCA Protein Assay Kit (Thermo Fisher Scientific, Cat# 23227). For each lysate, 30 µg of protein was mixed with an equal volume of 4x Laemmli sample buffer (Bio-Rad, Cat# 1610747) and the samples were boiled for 10 min at 75°C before loading onto a 4–15% Mini-PROTEAN TGX Precast Gel (Bio-Rad, Cat# 1610747). After electrophoresis, proteins were transferred onto a nitrocellulose membrane (Cytiva, Cat# 10600002) before primary and secondary antibody incubation and image acquisition using a Li-Cor Odyssey CLx Imager. Quantifications of relevant protein levels were performed using the Li-Cor ImageQuant Studio software. The primary antibodies used were rabbit anti-SYN3 (Synaptic Systems Cat#106303, RRID:AB_2619775; 1:2000) and mouse anti-VCL (MilliporeSigma Cat# V9131, RRID:AB_477629; 1:10000). The secondary antibodies used were: donkey anti-mouse IgG (H+L) DyLight 800 (Thermo Fisher Scientific Cat# SA5-10172, RRID:AB_2556752; 1:10,000) and donkey anti-rabbit IgG (H+L) DyLight 680 (Thermo Fisher Scientific Cat# SA5-10066, RRID:AB_2556646; 1:10,000).

#### Single-nucleus RNA sequencing and data analysis

Three *Cic^flox/flox^* and three *Emx1-Cre; Cic^flox/flox^* mice (littermates, two males and one female for each genotype) at P19 were euthanized with sodium pentobarbital and their hippocampus micro-dissected in ice-cold HBSS (Gibco, 14175103). The dissected tissues were snap-frozen in liquid nitrogen until nuclei extraction. One hippocampus from each mouse was used for nuclei extraction (i.e., three hippocampi per genotype). Nuclei were isolated using 10x Genomics Chromium Nuclei Isolation Kit (10x Genomics, Cat# 1000493) lysis buffer and the debris was removed using Miltenyi Anti-Nucleus beads (Miltenyi Biotec, Cat# 130-132-997). Briefly, frozen tissue pieces were pooled, weighed and chopped into small pieces on dry ice, before adding Lysis Reagent (10x Genomics, Cat# 2000558), Reducing Agent B (10x Genomics, Cat# 2000087), and Surfactant A (10x Genomics, Cat# 2000559). The tissue was homogenized using a glass douncer (MilliporeSigma, Cat# D9063) and lysed for 5 min on ice. A total of 500 µL of lysate was transferred to the Nuclei Isolation Column (10x Genomics, Cat# 2000562) to remove leftover tissue. The sample was then spun at 500 xg for 3 min at 4°C, and the nuclei pellet was resuspended in Nuclei Separation buffer composed of Nuclei Extraction buffer (Miltenyi Biotec, Cat# 130-128-024), PBS, 10% BSA and RNase Inhibitor (MilliporeSigma, Cat# 3335402001). Miltenyi anti-nucleus beads (Cat# 130-132-997) were then added to the sample (50 µL per million nuclei) and incubated for 15 min on ice followed by an additional 2 mL of the Nuclei Separation buffer. Then the sample was purified using an LS column (Miltenyi Biotec, Cat#130-042-401) attached to the MACS MultiStand (Miltenyi Biotec, Cat# 130-042-303) with a QuadroMACS separator (Miltenyi Biotec, Cat# 130-042-302). The purified nuclei were spun at 500 xg for 5 min, and the nuclei pellet was washed with wash buffer comprised of PBS, 10% BSA and RNase Inhibitor. The nuclei suspension was centrifuged at 4°C for 5 min at 500 xg again and the nuclei pellet was resuspended in wash buffer. Nuclei counting was done using a 1:1 dilution with acridine orange and propidium iodide on a haemocytometer (Thermo Fisher, Cat# 22-600-100). Following counting, the appropriate volume for each sample was calculated for a target capture of 20,000 nuclei and then the nuclei were loaded onto 10x 3’ GEM X chip. cDNA libraries were prepared as outlined by the Chromium GEM-X Single Cell 3’ Reagent Kits v4 user guide with modifications to the PCR cycles based on the calculated cDNA concentration. The molarity of each library was calculated based on library size as measured bioanalyzer (Agilent Technologies) and qPCR amplification data (Roche, Cat# 07960140001). Gene Expression libraries were sequenced on Illumina’s NovaSeqX with the following run parameters: read 1 – 28 cycles, read 2 – 90 cycles, index 1 – 10 cycles and index 2 – 10 cycles. The resulting FASTQ files were converted to count matrix files using Cell Ranger Count with Mouse (mm39) as the reference genome. Quality control, dimensionality reduction and initial clustering was performed in RStudio (2023.12.0 Build 369) using Seurat (v4.4.0; https://github.com/satijalab/seurat). Gene set enrichment analysis was performed using the Enrichr-KG(56) web server (https://maayanlab.cloud/enrichr-kg). Pseudotime analysis was performed using Monocle3(57–59). Ligand–receptor pairs analysis was performed using LIANA with the mouse consensus resource(41).

#### Confocal microscopy and image analyses

Immunofluorescent images were taken using a laser-scanning confocal microscope Zeiss LSM 700. For each animal, z-stacked images of the dentate gyrus and/or CA3 areas were acquired from at least two comparable coronal sections, and analyses were performed on images of these sections. Each data point represents the average value from multiple sections per animal. CALB1^+^ area measurements were carried out using ImageJ and were restricted to the CA3 stratum pyramidale region. The threshold of single fluorescence channel was adjusted, and particle analyses were then performed to yield the total area of CALB1^+^ mossy fiber terminals in the region of interest. The size of CALB1^+^ ZnT3^+^ MF terminals in CA3 stratum pyramidale was measured manually using ImageJ.

#### Statistics

Statistical analyses were performed using GraphPad Prism. Detailed information on statistical analysis and sample size is provided in the figures and their legends.

## Data availability

All data and materials for this manuscript are included in the Methods and Supplemental Information. Single-nucleus RNA-seq datasets have been deposited into GEO (GSE280854).

## Acknowledgements

We thank Dr. Sarah Hughes (University of Alberta, Edmonton, AB) for her assistance with confocal microscopy. We acknowledge the use of Princess Margaret Genomics Centre at the University Health Networks for single-nucleus RNA-seq services.

## Conflict of interest statement

The authors have declared that no conflict of interest exists.

## Funding

Qiumin Tan receives support from the Natural Sciences and Engineering Research Council of Canada (RGPIN□2019□06153) and the Canada Foundation for Innovation (Award 38985). This work was supported by a grant from *The Scottish Rite Charitable Foundation of Canada*. Qiumin Tan is a Tier 2 Canada Research Chair in Neurodevelopmental Disorders. This study was undertaken, in part, thanks to funding from the Canada Research Chairs program.

## Author contributions statement

Q.T. conceptualized the project and designed experiments. All authors performed research and analyzed data. All authors wrote and edited the paper.

## Notes

### Competing Interest Statement

The authors have declared no competing interest.

## References

1 Kolodkin, A.L. and Tessier-Lavigne, M. (2011) Mechanisms and molecules of neuronal wiring: a primer. Cold Spring Harb Perspect Biol, 3.

2 Agi, E., Kulkarni, A. and Hiesinger, P.R. (2020) Neuronal strategies for meeting the right partner during brain wiring. Current Opinion in Neurobiology, 63, 1–8.

3 Stoeckli, E.T. (2018) Understanding axon guidance: are we nearly there yet? Development, 145.

4 Van Battum, E.Y., Gunput, R.A., Lemstra, S., Groen, E.J., Yu, K.L., Adolfs, Y., Zhou, Y., Hoogenraad, C.C., Yoshida, Y., Schachner, M., et al. (2014) The intracellular redox protein MICAL-1 regulates the development of hippocampal mossy fibre connections. Nat Commun, 5, 4317.

5 Evstratova, A. and Toth, K. (2014) Information processing and synaptic plasticity at hippocampal mossy fiber terminals. Front Cell Neurosci, 8, 28.

6 Suto, F., Tsuboi, M., Kamiya, H., Mizuno, H., Kiyama, Y., Komai, S., Shimizu, M., Sanbo, M., Yagi, T., Hiromi, Y. et al. (2007) Interactions between plexin-A2, plexin-A4, and semaphorin 6A control lamina-restricted projection of hippocampal mossy fibers. Neuron, 53, 535–547.

7 Seki, T. and Rutishauser, U. (1998) Removal of polysialic acid-neural cell adhesion molecule induces aberrant mossy fiber innervation and ectopic synaptogenesis in the hippocampus. J Neurosci, 18, 3757–3766.

8 Amaral, D.G. and Dent, J.A. (1981) Development of the mossy fibers of the dentate gyrus: I. A light and electron microscopic study of the mossy fibers and their expansions. J Comp Neurol, 195, 51–86.

9 Tawarayama, H., Yoshida, Y., Suto, F., Mitchell, K.J. and Fujisawa, H. (2010) Roles of semaphorin-6B and plexin-A2 in lamina-restricted projection of hippocampal mossy fibers. J Neurosci, 30, 7049–7060.

10 Zhao, X.F., Kohen, R., Parent, R., Duan, Y., Fisher, G.L., Korn, M.J., Ji, L., Wan, G., Jin, J., Puschel, A.W. et al. (2018) PlexinA2 Forward Signaling through Rap1 GTPases Regulates Dentate Gyrus Development and Schizophrenia-like Behaviors. Cell Rep, 22, 456–470.

11 Toda, T., Parylak, S.L., Linker, S.B. and Gage, F.H. (2019) The role of adult hippocampal neurogenesis in brain health and disease. Mol Psychiatry, 24, 67–87.

12 Kempermann, G. (2022) What Is Adult Hippocampal Neurogenesis Good for? Front Neurosci, 16, 852680.

13 Lipp, H.P., Schwegler, H. and Driscoll, P. (1984) Postnatal modification of hippocampal circuitry alters avoidance learning in adult rats. Science, 225, 80–82.

14 Parent, J.M., Yu, T.W., Leibowitz, R.T., Geschwind, D.H., Sloviter, R.S. and Lowenstein, D.H. (1997) Dentate granule cell neurogenesis is increased by seizures and contributes to aberrant network reorganization in the adult rat hippocampus. J Neurosci, 17, 3727–3738.

15 Lu, H.C., Tan, Q., Rousseaux, M.W., Wang, W., Kim, J.Y., Richman, R., Wan, Y.W., Yeh, S.Y., Patel, J.M., Liu, X. et al. (2017) Disruption of the ATXN1-CIC complex causes a spectrum of neurobehavioral phenotypes in mice and humans. Nat Genet, 49, 527–536.

16 Hwang, I., Pan, H., Yao, J., Elemento, O., Zheng, H. and Paik, J. (2020) CIC is a critical regulator of neuronal differentiation. JCI Insight, in press.

17 Ahmad, S.T., Rogers, A.D., Chen, M.J., Dixit, R., Adnani, L., Frankiw, L.S., Lawn, S.O., Blough, M.D., Alshehri, M., Wu, W. et al. (2019) Capicua regulates neural stem cell proliferation and lineage specification through control of Ets factors. Nat Commun, 10, 2000.

18 Tan, Q. and Zoghbi, H.Y. (2019) Mouse models as a tool for discovering new neurological diseases. Neurobiology of Learning and Memory, 165, 106902.

19 Kishnani, S., Riley, K., Mikati, M.A. and Jiang, Y.-h. (2020) Phenotypic Variability of an Inherited Pathogenic Variant in CIC Gene: A New Case Report in Two-Generation Family and Literature Review. Journal of Pediatric Neurology, in press.

20 Cao, X., Wolf, A., Kim, S.E., Cabrera, R.M., Wlodarczyk, B.J., Zhu, H., Parker, M., Lin, Y., Steele, J.W., Han, X., et al. (2020) CIC de novo loss of function variants contribute to cerebral folate deficiency by downregulating FOLR1 expression. J Med Genet, in press.

21 Vissers, L.E., de Ligt, J., Gilissen, C., Janssen, I., Steehouwer, M., de Vries, P., van Lier, B., Arts, P., Wieskamp, N., del Rosario, M., et al. (2010) A de novo paradigm for mental retardation. Nat Genet, 42, 1109–1112.

22 Sharma, S., Hourigan, B., Patel, Z., Rosenfeld, J.A., Chan, K.M., Wangler, M.F., Yi, J.S., Lehman, A., Study, C., Horvath, G. et al. (2022) Novel CIC variants identified in individuals with neurodevelopmental phenotypes. Hum Mutat, 43, 889–899.

23 Ruiz, I., Wiltrout, K., Stredny, C. and Mahida, S. (2024) CIC-Related Neurodevelopmental Disorder: A Review of the Literature and an Expansion of Genotype and Phenotype. Genes (Basel), 15.

24 Almutair, M., Thabet, F., Hundallah, K. and Tabarki, B. (2024) CIC variants and folinic acid-responsive seizures. Mol Genet Metab, 143, 108574.

25 Hwang, I., Pan, H., Yao, J., Elemento, O., Zheng, H. and Paik, J. (2020) CIC is a critical regulator of neuronal differentiation. JCI Insight, 5.

26 Hourigan, B., Balay, S.D., Yee, G., Sharma, S. and Tan, Q. (2021) Capicua regulates the development of adult-born neurons in the hippocampus. Sci Rep, 11, 11725.

27 Tian, C., Gong, Y., Yang, Y., Shen, W., Wang, K., Liu, J., Xu, B., Zhao, J. and Zhao, C. (2012) Foxg1 has an essential role in postnatal development of the dentate gyrus. J Neurosci, 32, 2931–2949.

28 Gorski, J.A., Talley, T., Qiu, M., Puelles, L., Rubenstein, J.L. and Jones, K.R. (2002) Cortical excitatory neurons and glia, but not GABAergic neurons, are produced in the Emx1-expressing lineage. J Neurosci, 22, 6309–6314.

29 Liu, X.D., Zhu, X.N., Halford, M.M., Xu, T.L., Henkemeyer, M. and Xu, N.J. (2018) Retrograde regulation of mossy fiber axon targeting and terminal maturation via postsynaptic Lnx1. J Cell Biol, 217, 4007–4024.

30 Wenzel, H.J., Cole, T.B., Born, D.E., Schwartzkroin, P.A. and Palmiter, R.D. (1997) Ultrastructural localization of zinc transporter-3 (ZnT-3) to synaptic vesicle membranes within mossy fiber boutons in the hippocampus of mouse and monkey. Proc Natl Acad Sci U S A, 94, 12676–12681.

31 Sindreu, C., Palmiter, R.D. and Storm, D.R. (2011) Zinc transporter ZnT-3 regulates presynaptic Erk1/2 signaling and hippocampus-dependent memory. Proc Natl Acad Sci U S A, 108, 3366–3370.

32 Apostolo, N., Smukowski, S.N., Vanderlinden, J., Condomitti, G., Rybakin, V., Ten Bos, J., Trobiani, L., Portegies, S., Vennekens, K.M., Gounko, N.V. et al. (2020) Synapse type-specific proteomic dissection identifies IgSF8 as a hippocampal CA3 microcircuit organizer. Nat Commun, 11, 5171.

33 Pieribone, V.A., Porton, B., Rendon, B., Feng, J., Greengard, P. and Kao, H.T. (2002) Expression of synapsin III in nerve terminals and neurogenic regions of the adult brain. J Comp Neurol, 454, 105–114.

34 Gerfen, C.R., Paletzki, R. and Heintz, N. (2013) GENSAT BAC cre-recombinase driver lines to study the functional organization of cerebral cortical and basal ganglia circuits. Neuron, 80, 1368–1383.

35 Nakazawa, K., Quirk, M.C., Chitwood, R.A., Watanabe, M., Yeckel, M.F., Sun, L.D., Kato, A., Carr, C.A., Johnston, D., Wilson, M.A. et al. (2002) Requirement for hippocampal CA3 NMDA receptors in associative memory recall. Science, 297, 211–218.

36 Ubina, T., Vahedi-Hunter, T., Agnew-Svoboda, W., Wong, W., Gupta, A., Santhakumar, V. and Riccomagno, M.M. (2021) ExBoX - a simple Boolean exclusion strategy to drive expression in neurons. J Cell Sci, 134.

37 Kim, J.Y., Grunke, S.D., Levites, Y., Golde, T.E. and Jankowsky, J.L. (2014) Intracerebroventricular viral injection of the neonatal mouse brain for persistent and widespread neuronal transduction. J Vis Exp, in press., 51863.

38 Kim, J.Y., Ash, R.T., Ceballos-Diaz, C., Levites, Y., Golde, T.E., Smirnakis, S.M. and Jankowsky, J.L. (2013) Viral transduction of the neonatal brain delivers controllable genetic mosaicism for visualising and manipulating neuronal circuits in vivo. Eur J Neurosci, 37, 1203–1220.

39 Hochgerner, H., Zeisel, A., Lonnerberg, P. and Linnarsson, S. (2018) Conserved properties of dentate gyrus neurogenesis across postnatal development revealed by single-cell RNA sequencing. Nat Neurosci, 21, 290–299.

40 Lee, Y. (2020) Regulation and function of capicua in mammals. Exp Mol Med, 52, 531–537.

41 Dimitrov, D., Turei, D., Garrido-Rodriguez, M., Burmedi, P.L., Nagai, J.S., Boys, C., Ramirez Flores, R.O., Kim, H., Szalai, B., Costa, I.G. et al. (2022) Comparison of methods and resources for cell-cell communication inference from single-cell RNA-Seq data. Nat Commun, 13, 3224.

42 Kolodkin, A.L., Levengood, D.V., Rowe, E.G., Tai, Y.T., Giger, R.J. and Ginty, D.D. (1997) Neuropilin is a semaphorin III receptor. Cell, 90, 753–762.

43 He, Z. and Tessier-Lavigne, M. (1997) Neuropilin is a receptor for the axonal chemorepellent Semaphorin III. Cell, 90, 739–751.

44 Takahashi, T., Fournier, A., Nakamura, F., Wang, L.H., Murakami, Y., Kalb, R.G., Fujisawa, H. and Strittmatter, S.M. (1999) Plexin-neuropilin-1 complexes form functional semaphorin-3A receptors. Cell, 99, 59–69.

45 Galimberti, I., Gogolla, N., Alberi, S., Santos, A.F., Muller, D. and Caroni, P. (2006) Long-term rearrangements of hippocampal mossy fiber terminal connectivity in the adult regulated by experience. Neuron, 50, 749–763.

46 Seeger, M.A. and Beattie, C.E. (1999) Attraction versus repulsion: modular receptors make the difference in axon guidance. Cell, 97, 821–824.

47 Evans, T.A. and Bashaw, G.J. (2010) Axon guidance at the midline: of mice and flies. Curr Opin Neurobiol, 20, 79–85.

48 Castellani, V. (2013) Building Spinal and Brain Commissures: Axon Guidance at the Midline. International Scholarly Research Notices, 2013, 315387.

49 Van Battum, E.Y., Brignani, S. and Pasterkamp, R.J. (2015) Axon guidance proteins in neurological disorders. Lancet Neurol, 14, 532–546.

50 Verhagen, M.G. and Pasterkamp, R.J. (2020) Rubenstein, J., Rakic, P., Chen, B., Kwan, K.Y., Kolodkin, A. and Anton, E. (eds.), In Cellular Migration and Formation of Axons and Dendrites (Second Edition). Academic Press, in press., pp. 109–122.

51 Weth, F. and Kania, A. (2020) Rubenstein, J., Rakic, P., Chen, B., Kwan, K.Y., Kolodkin, A. and Anton, E. (eds.), In Cellular Migration and Formation of Axons and Dendrites (Second Edition). Academic Press, in press., pp. 123–146.

52 Nishimura-Akiyoshi, S., Niimi, K., Nakashiba, T. and Itohara, S. (2007) Axonal netrin-Gs transneuronally determine lamina-specific subdendritic segments. Proc Natl Acad Sci U S A, 104, 14801–14806.

53 Tan, Q. and Zoghbi, H.Y. (2019) Mouse models as a tool for discovering new neurological diseases. Neurobiol Learn Mem, 165, 106902.

54 Kishnani, S., Riley, K., Mikati, M.A. and Jiang, Y.-h. (2020) Phenotypic Variability of an Inherited Pathogenic Variant in CIC Gene: A New Case Report in Two-Generation Family and Literature Review. Journal of Pediatric Neurology, 19, 193–201.

55 van Bruggen, R., Patel, Z.H., Wang, M., Suk, T.R., Rousseaux, M.W.C. and Tan, Q. (2023) A Versatile Strategy for Genetic Manipulation of Cajal-Retzius Cells in the Adult Mouse Hippocampus. eNeuro, in press.

56 Evangelista, J.E., Xie, Z., Marino, G.B., Nguyen, N., Clarke, D.J.B. and Ma’ayan, A. (2023) Enrichr-KG: bridging enrichment analysis across multiple libraries. Nucleic Acids Res, 51, W168–W179.

57 Qiu, X., Mao, Q., Tang, Y., Wang, L., Chawla, R., Pliner, H.A. and Trapnell, C. (2017) Reversed graph embedding resolves complex single-cell trajectories. Nat Methods, 14, 979–982.

58 Trapnell, C., Cacchiarelli, D., Grimsby, J., Pokharel, P., Li, S., Morse, M., Lennon, N.J., Livak, K.J., Mikkelsen, T.S. and Rinn, J.L. (2014) The dynamics and regulators of cell fate decisions are revealed by pseudotemporal ordering of single cells. Nat Biotechnol, 32, 381–386.

59 Cao, J., Spielmann, M., Qiu, X., Huang, X., Ibrahim, D.M., Hill, A.J., Zhang, F., Mundlos, S., Christiansen, L., Steemers, F.J. et al. (2019) The single-cell transcriptional landscape of mammalian organogenesis. Nature, 566, 496–502.

