## Supplemental information for "Capicua refines mossy fiber–CA3 axon targeting in the late postnatal hippocampus"

**Figure S1**

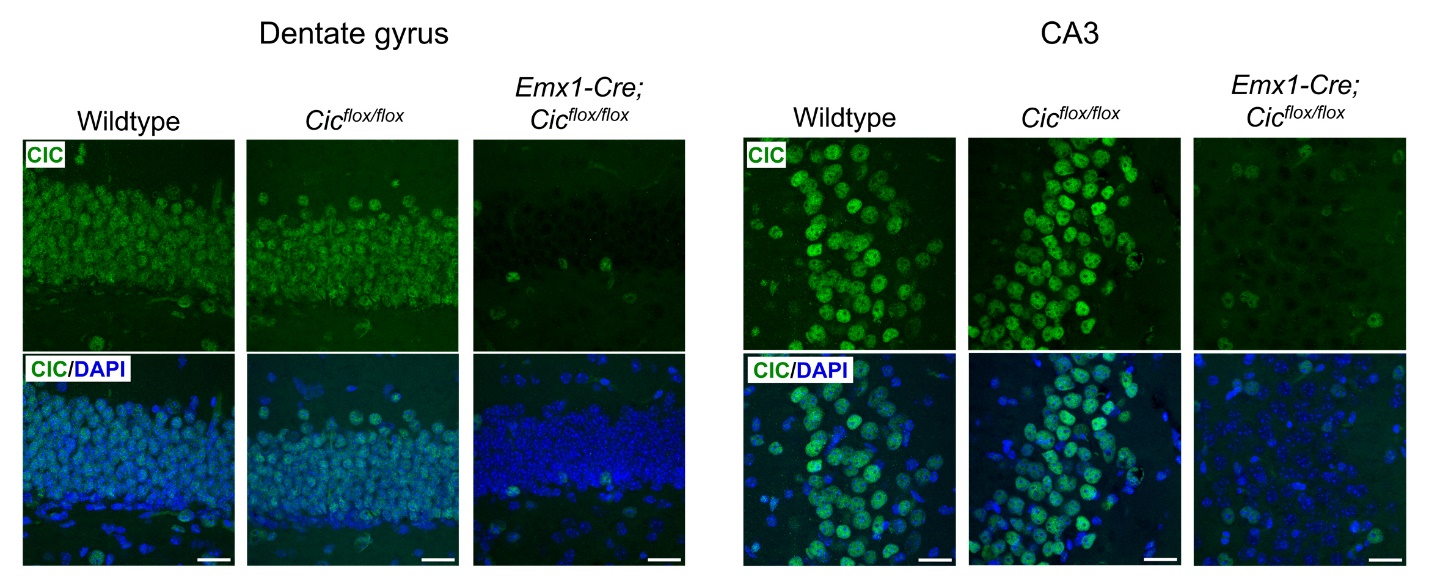

**Figure S1. Efficient CIC deletion in dentate gyrus granule neurons and CA3 neurons of *Emx1-Cre; Cic^flox/flox^* knockout mice.** Immunofluorescence staining for CIC in 8-week-old animals. Wildtype and *Cic^flox/flox^* mice show comparable CIC immunoreactivity. There is a marked loss of CIC signal in most dentate gyrus and CA3 neurons of knockout mice. CIC remains detectable in non-*Emx1*-lineage cells, such as inhibitory neurons and microglia. CA1, cornu ammonis 1; CA3, cornu ammonis 3; DG, dentate gyrus. Scale bars = 25 µm.

**Figure S2**

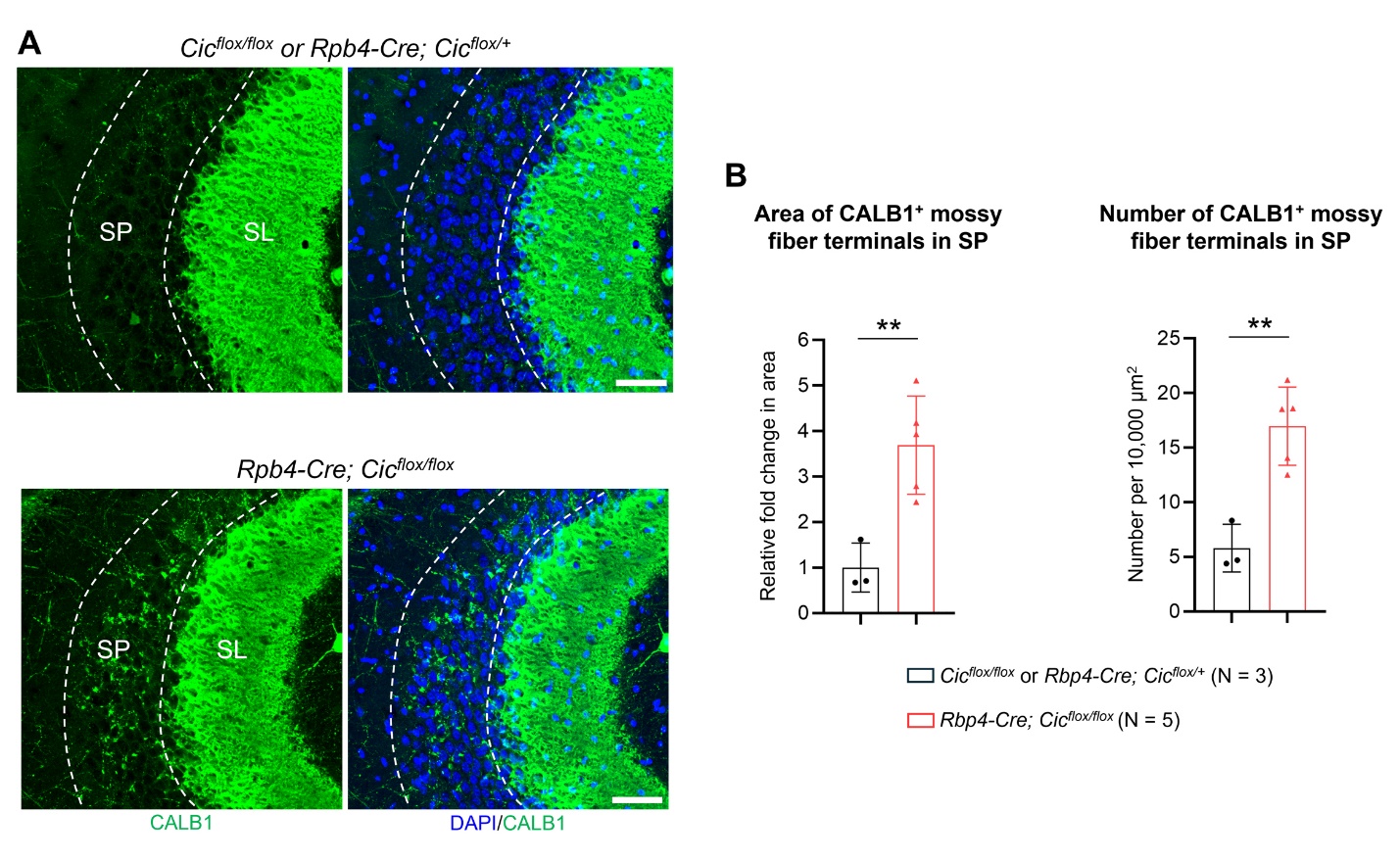

**Figure S2. CIC is required in dentate gyrus granule neurons for lamina-specific mossy fiber innervation.** (**A**) Confocal images of 8-week-old control (*Cic^flox/flox^* or *Rbp4-Cre; Cic^flox/+^*) and *Rbp4-Cre; Cic^flox/flox^* knockout mice. In *Rbp4-Cre; Cic^flox/flox^* mice, CALB1^+^ mossy fiber terminals abnormally innervate the stratum pyramidale (SP) in the CA3a region. SL, stratum lucidum. Scale bars = 50 µm. (**B**) Quantification of CALB^+^ MF terminal area and number in the SP. Data are presented as scatter plots, with error bars indicating mean ± SD. Statistical analysis was performed using Welch’s t test. **, *P* < 0.01.

**Figure S3**

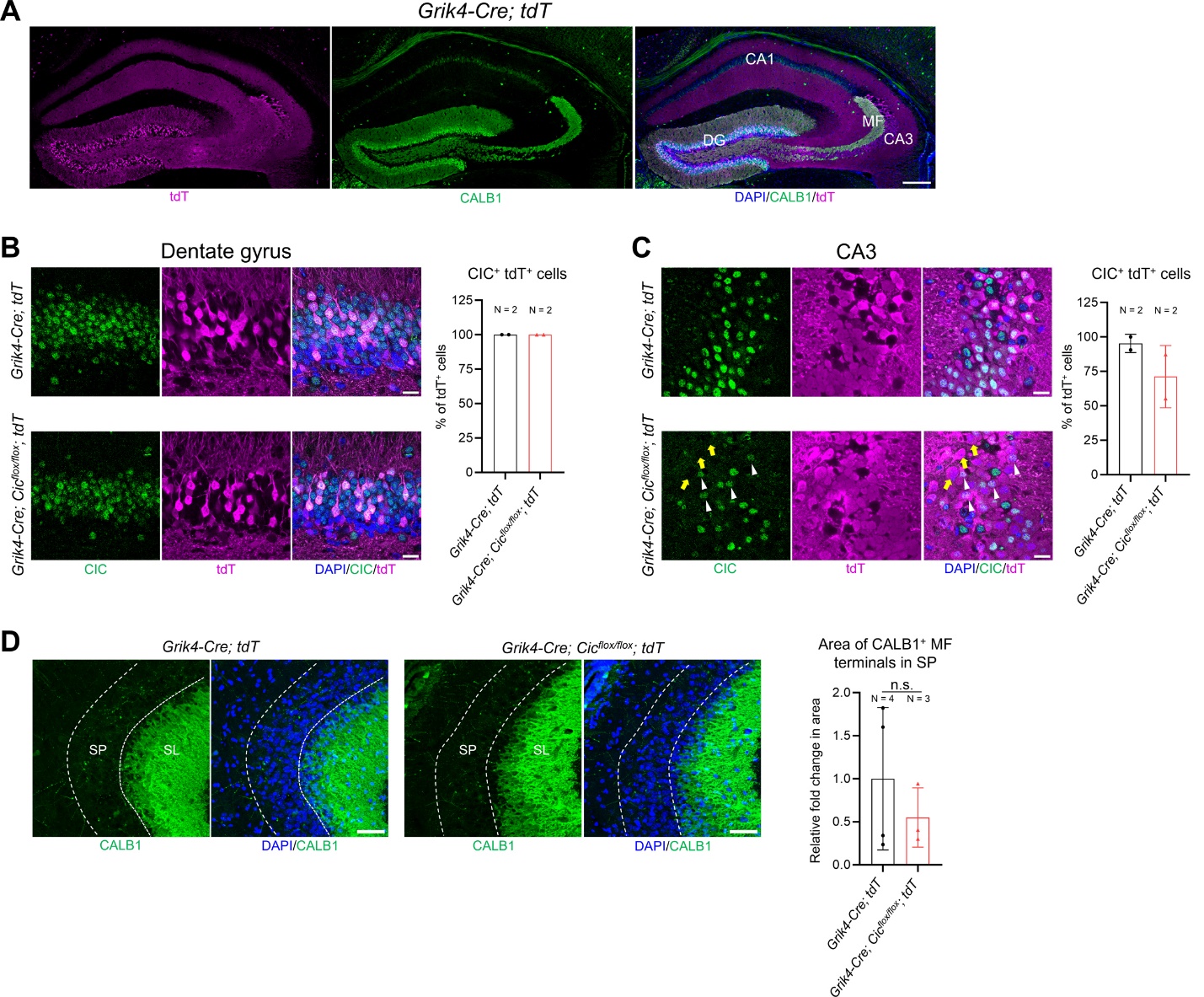

**Figure S3. Partial loss of CIC in CA3 neurons does not affect mossy fiber projections.** (**A**) Confocal images of the hippocampus from *Grik4-Cre; tdT* mice at postnatal day (P) 20, showing Cre-mediated recombination in dentate gyrus (DG) and CA3 region. Scale bar = 250 µm. CA1, cornu ammonis 1; CA3, cornu ammonis 3; MF, mossy fiber. (**B**) Immunostaining for CIC and tdTomato (tdT) reveals the presence of CIC immunoreactivity in tdT^+^ granule neurons in both genotypes. This indicates that, while the *Grik4-Cre* driver efficiently recombines the *tdT* reporter allele in granule neurons, it does not mediate recombination of the *Cic^flox^* allele in these cells. Quantification of the proportion of tdT^+^ granule neurons expressing CIC is shown on the right. Scale bars = 20 µm. (**C)** Confocal images of CIC and tdTomato (tdT) immunostaining show that the *Grik4-Cre* driver mediates recombination of the *Cic^flox^* allele in a subset of CA3 neurons. White arrowheads indicate tdT^+^ neurons expressing CIC, while yellow arrows indicate tdT^+^ neurons lacking CIC expression. Quantification of the proportion of tdT^+^ CA3 neurons expressing CIC is shown on the right. Scale bars = 20 µm. (**D**) Normal MF innervation in CA3 stratum pyramidale (SP) of *Grik4-Cre; Cic^flox/flox^; tdT* mice at P20. Quantification of CALB^+^ MF terminal area in the SP is shown on the right. Scale bars = 50 µm. Data are presented as scatter plots with error bars representing mean ± SD. Statistical analysis was performed using Welch’s t test. n.s., not significant.

**Figure S4**

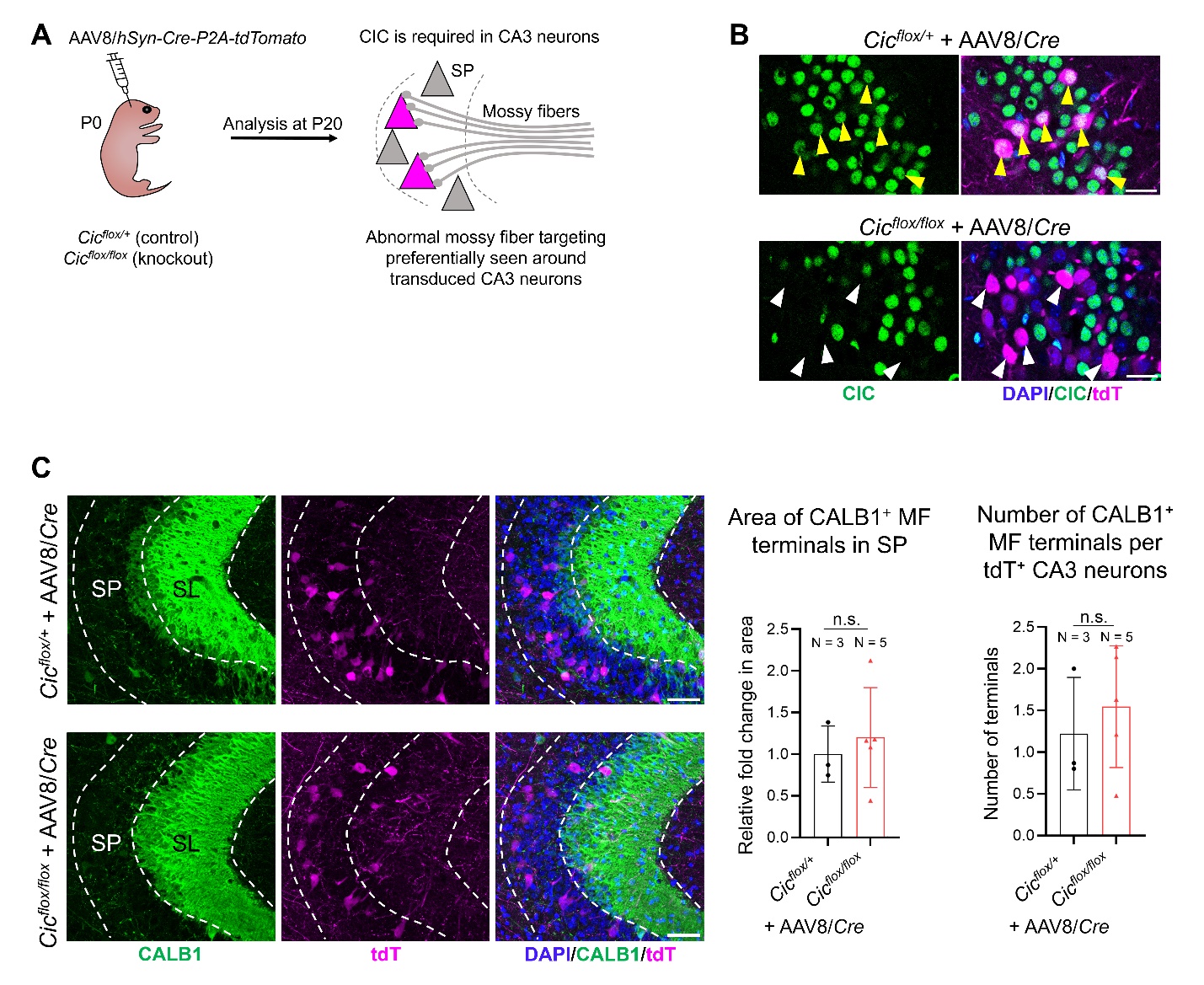

**Figure S4. Adeno associated virus (AAV)-mediated genetic mosaicism for studies of cell-autonomy.** (**A**) A schematic diagram outlines the experimental approach. Intraventricular injection of AAV to postnatal day (P) 0 pups is used to deliver the bicistronic Cre recombinase and tdTomato (tdT) reporter under the neuronal human synapsin promoter (*hSyn*). AAV-injected animals were analyzed at P20. (**B**) Representative confocal images demonstrate that CA3 neurons are sparsely transduced. CIC remains expressed in AAV8/*Cre*-transduced CA3 neurons in the *Cic^flox/+^* mice (yellow arrowheads), but it is efficiently deleted from AAV8/*Cre*-transduced neurons in the *Cic^flox/flox^* mice (white arrowheads). Scale bars = 25 µm. (**C**) Confocal images of immunostaining for CALB1 and tdT in the CA3a region. Note the lack of strong tdT signal in the stratum lucidum (SL), indicating that granule neurons are minimally transduced in this system. Mosaic ablation of CIC from CA3 neurons does not cause wildtype (non-transduced) mossy fiber (MF) to abnormally innervate the cell bodies of CIC-deleted (tdT^+^, transduced) CA3 neurons. Quantification of the CALB1^+^ MF terminal area and number in CA3 SP is shown on the right. Data are presented in scatter plots with error bars representing ± SD. Statistical analysis was performed using Welch’s t test. n.s., not significant.

**Figure S5**

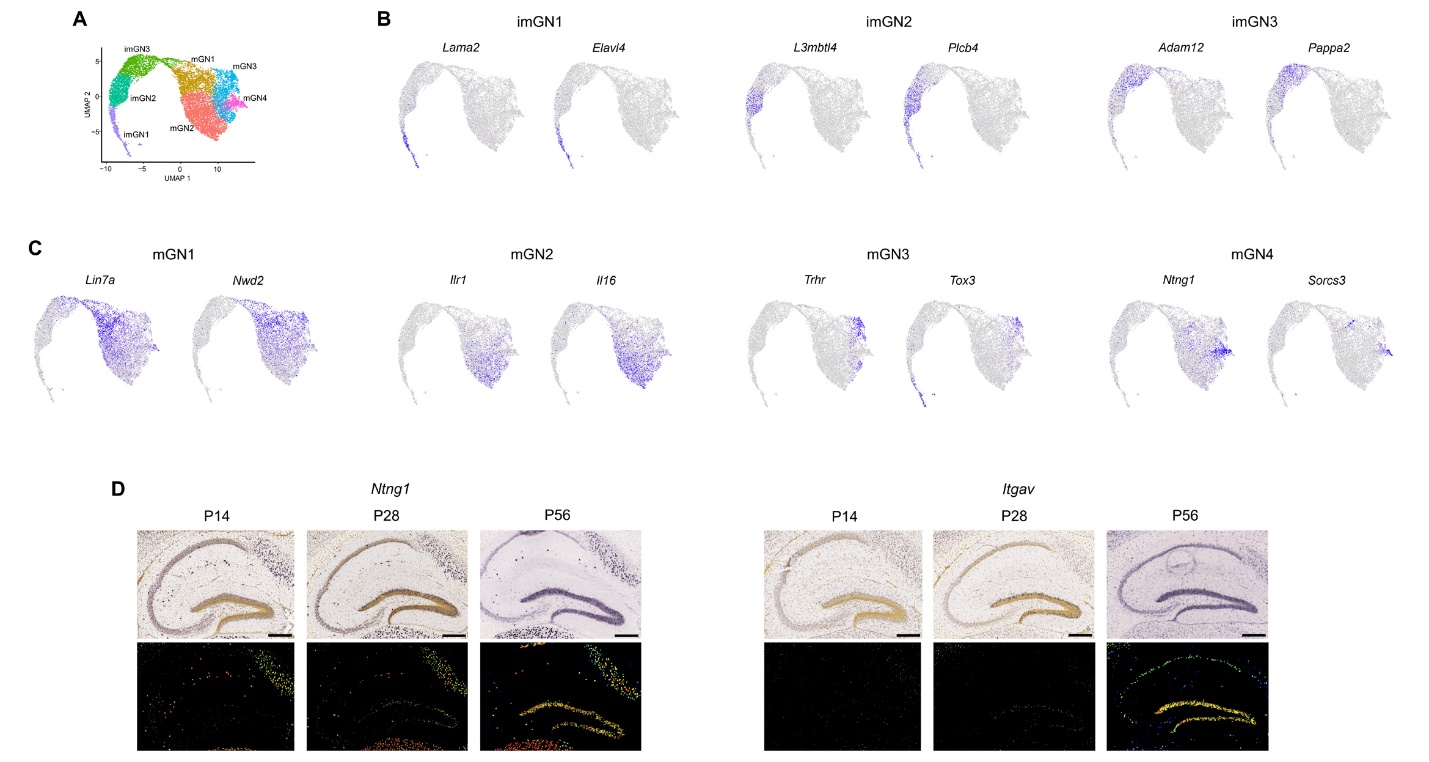

**Figure S5. Marker analysis of granule neuron subsets.** (**A**) UMAP plot showing seven clusters of granule neurons, including immature granule neurons (imGN) and mature granule neurons (mGN). (**B**, **C**) UMAP feature plots illustrating the expression of cluster-specific markers for immature (**B**) and mature (**C**) granule neuron clusters. (**D**) Expression patterns of the mature granule neuron (mGN4) makers *Ntng1* and *Itgav* in the mouse hippocampus at postnatal day (P) 14, P28, and P56. In situ hybridization images are shown at the top and color-coded expression maps are shown at the bottom. At P14, both genes are expressed at low levels. By P28, granule neurons in the outer granular layers, indicative of mature granule neurons, start to express *Ntng1* and *Itgav*. At P56, both genes are highly expressed throughout the granular layer, consistent with mature granule neuron localization. Data from Allen Mouse Brain Atlas. Scale bars = 300 µm.

**Figure S6**

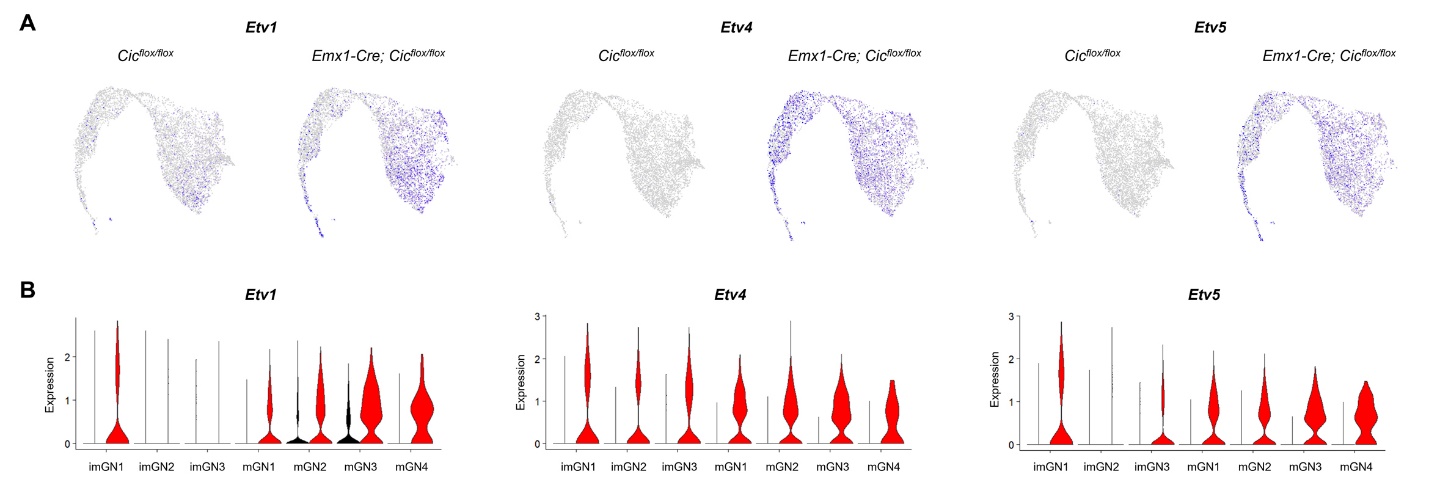

**Figure S6. Expression of CIC target genes (*Etv1*, *Etv4*, and *Etv5*) across cell clusters.** (**A**) UMAP feature plots showing the expression patterns of CIC target genes between control and knockout genotypes. (**B**) Violin plots showing upregulation of CIC target genes in cell clusters from *Emx1-Cre; Cic^flox/flox^* knockout mice.
